# Nanopore genome skimming with Illumina polishing yields highly accurate mitogenome sequences: a case study of *Niphargus* amphipods

**DOI:** 10.1101/2025.09.24.678374

**Authors:** Alice Salussolia, Ninon Lecoquierre, Fabio Stoch, Jean-François Flot

## Abstract

With over 430 species currently described, the amphipod genus *Niphargus* Schiödte, 1849 is the most species-rich crustacean genus in subterranean waters. Previous phylogenetic studies of this genus have relied mainly on mitochondrial COI and nuclear 28S sequences, which do not resolve all the nodes in its phylogeny. As a first step towards a mitogenome-based phylogeny of niphargids, we present here the first complete mitogenome sequence of *Niphargus*. To obtain high-accuracy mitogenome sequences and annotations, genome skimming of three individuals of *Niphargus dolenianensis* Lorenzi, 1898 was performed using both short, accurate reads (Illumina) and long, noisier reads (nanopore). Whereas the direct assembly of Illumina sequences yielded structurally incorrect mitogenome sequences, the assembly of nanopore reads produced highly accurate sequences that were corroborated by the mapping of Illumina reads. Polishing the nanopore consensus using Illumina reads corrected a handful of errors at the homopolymer level. The resulting mitogenome sequences ranged from 14,956 to 15,199 bp and shared the same arrangement of 13 protein-coding genes, two ribosomal RNA genes, 22 transfer RNA genes, and a putative control region. Phylogenetic analyses based on protein-coding genes confirmed that the Niphargidae family is sister to Pseudoniphargidae, resolving their relationships with other amphipod families. This highlights the utility of mtDNA genome sequences for studying the evolution of this groundwater genus, and the refinement of new methodological approaches, such as nanopore sequencing, is promising for the study of its origin and diversification.

## Introduction

Groundwater constitutes an extensive and globally prevalent ecosystem that contains the majority of accessible freshwater resources (Saccò et al., 2024). Animals inhabiting groundwater play a crucial role in enhancing the biodiversity of freshwater ecosystems and are essential for processes such as nutrient cycling and bioturbation (Bardgett and Van Der Putten, 2014). Groundwater offers numerous ecosystem services, ranging from supporting terrestrial and surface freshwater ecosystems to supplying drinking water (Griebler and Avramov, 2015). Therefore, investigating groundwater animals is vital for advancing our understanding of groundwater community structure and processes, ecosystem services, and their reactions to environmental changes (Maurice and Bloomfield, 2012).

With a remarkable diversity of over 430 identified species (Horton et al., 2023), the amphipod genus *Niphargus* Schiödte, 1849 is the most species-rich crustacean genus in subterranean waters. The distribution of this genus spans the Iberian Peninsula to the west and Iran to the east, covering European regions south of the limits of Quaternary glaciations (Zagmajster et al., 2014). Studies on *Niphargus* are hampered by the lack of phylogenetic information on most species (except in a few well-studied areas) (Zagmajster et al., 2014) and by the frequent occurrence of complexes of cryptic and pseudocryptic species *Niphargus* (Stoch et al., 2022). To date, studies on the phylogeny of the genus *Niphargus* have relied mainly on mitochondrial COI and nuclear 28S rRNA marker genes, and the resulting phylogenies remain poorly resolved, particularly for deeper nodes (Stoch et al., 2024b). Complete mitochondrial genome sequences of animals have been reported to yield well-resolved phylogenies (Cameron et al., 2007), but so far, not a single complete mitogenome sequence of *Niphargus* has been published.

To fill this gap and begin investigating the potential of niphargid complete mitochondrial genome sequences to shed new light on amphipod phylogenetic relationships, we focused on the Alpine species *Niphargus dolenianensis* Lorenzi, 1898 and used genome skimming (Dodsworth, 2015) to assemble its complete mitogenome. Although the family Niphargidae is very diverse (over 230 described species following Horton et al., 2023), most species belong to the so-called megaclade (Stoch et al., 2024a). Therefore, we selected one widely distributed species from the southern Alps (northern Italy) and a part of this megaclade for our study. We aimed to obtain a highly reliable *Niphargus dolenianensis* mitogenome for future comparative studies. For this purpose, we sequenced three different individuals using both nanopore and Illumina platforms and used Illumina reads to polish the assembly of nanopore reads. Finally, to propose guidelines for future studies targeting niphargid mitogenomes, we tested whether Illumina-only and nanopore-only genome skimming would have been sufficient to obtain high-quality mitogenome sequences for downstream analysis.

## Material and methods

### DNA extraction and sequencing

Specimens of *Niphargus dolenianensis* Lorenzi, 1898 (Fig.1) were collected in 2024 from two springs and a brook using a hand-net. Information on the specimens used in the analyses, collection sites, and DNA vouchers is presented in Table 1. Fresh samples were immediately preserved in 96% EtOH and then stored at -20°C at the Evolutionary Biology & Ecology unit of the Université libre de Bruxelles (ULB), Belgium. DNA extraction was performed from one or two pereopods (depending on the size of the specimen) using the Macherey-Nagel NucleoSpin Tissue kit, following the manufacturer’s protocol. The eluted DNAs was stored at -20°C. The three DNA extracts were sent for Illumina sequencing at BRIGHTcore facility (Brussels, Belgium), enzymatic fragmentation was performed (2 x 100 bp paired-end reads) following by PCR-free library preparation, run on Illumina NovaSeq 6000 machine. The same three DNA extracts were also used for long-read nanopore sequencing using Oxford Nanopore Technologies (ONT) with the Rapid PCR Barcoding kit SQK-RPB114.24, on a PromethION R10.4.1 flow cell.

**Figure 1.**
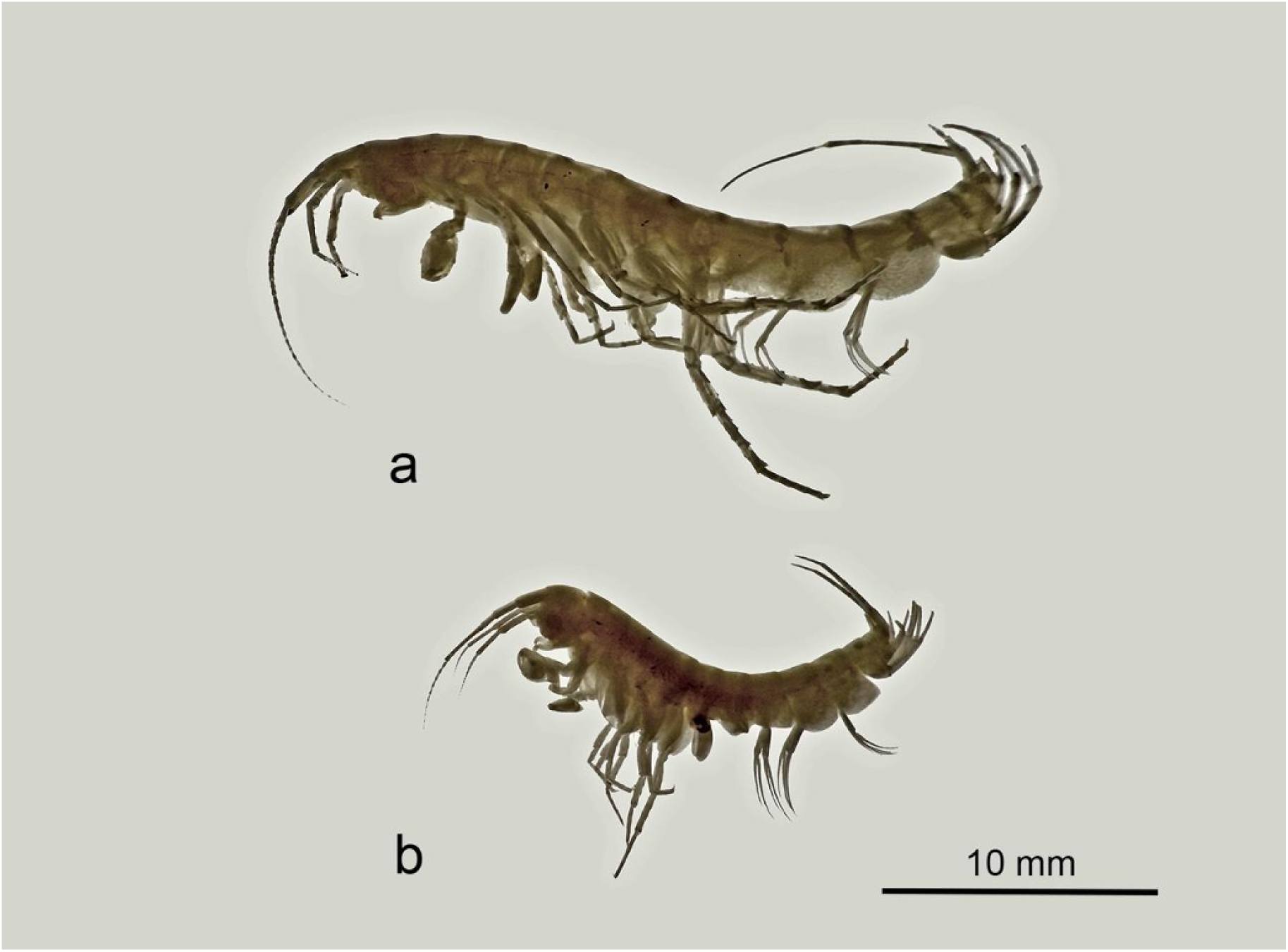
*Niphargus dolenianensis*, habitus (a): male; (b): female

**Table 1.** List of specimens used in the study with the GenBank accession numbers of the deposited mitogenome sequences. All specimens were collected on 08/04/2024 by F. Stoch and G. Tomasin then identified morphologically.

| Voucher | Accession number | Locality | Latitude (WGS84) | Longitude (WGS 84) | Altitude (m a.s.l.) |
| --- | --- | --- | --- | --- | --- |
| FS_24.009 | PV534572 | San Giovanni al Natisone, spring Villa Trento, Dolegnano, Italy | 13.420386 | 45.986732 | 80 |
| FS_24.010 | PV534571 | Gorizia, spring along Vallone dell'Acqua road, Italy | 13.597421 | 45.954605 | 163 |
| FS_24.011 | PV534570 | Gorizia, Cormons, Forest of Plessiva, springbrook, Italy | 13.488482 | 45.977869 | 97 |

The Illumina sequencing run cost us €3.90 per million reads, and from the time the DNA samples reached the laboratory, the turnaround time for obtaining raw data was four weeks (costs and processing times may vary among providers). For nanopore sequencing, the SQK-RPB114.24 kit (https://store.nanoporetech.com/rapid-pcr-barcoding-kit-24-v14.html) costs roughly €680 and supports 24 barcoded samples, with the kit usable for up to six runs, resulting in an estimated cost of €4.70 per sample. Preparation of the nanopore library required 15 min +PCR, and raw sequencing data were available within 24 h.

### Mitochondrial genome annotation and analysis

Illumina data were trimmed using Skewer (Jiang et al., 2014) in paired-end mode. Nanopore data were basecalled using Guppy v3.8 (in super-high-quality mode) to generate fastq reads from the raw fast5. Nanopore reads were filtered to retain only reads with an average quality score of at least 14 using the script split_qscore.py available as part of the Buttery-eel package (Samarakoon et al., 2023) and assembled using Flye v2.9.6 (Kolmogorov et al., 2019), as well as with hifiasm v0.25.0-r726 (Cheng et al., 2021) for comparison. The resulting graphical fragment assembly (GFA) files were examined using Bandage (Wick et al., 2015) to identify and extract the circular contig of the mitochondrial genome. The first automatic Illumina polishing of the nanopore-assembled mitochondrial contig was performed using Polypolish (Wick and Holt, 2022). The resulting assemblies were checked closely for structural and base-level errors by mapping the nanopore and Illumina reads on them using minimap2 (Li, 2018), converting the resulting SAM file into BAM, sorting it using SAMtools (Li et al., 2009), and visualizing the resulting read pileups using Tablet (Milne et al., 2010). The few remaining errors detected in Tablet were corrected manually in the contig FASTA files using AliView (Larsson, 2014). For comparison, Illumina-only assemblies of the three mitochondrial genomes were attempted using the NOVO-plasty assembler (Dierckxsens et al., 2017), using as seed the previously published COI sequence (Genbank accession KY706720) of a *Niphargus dolenianensis* individual (Eme et al., 2018).

A quick annotation of each nanopore-assembled, Illumina-polished mitogenome sequence was performed using GeSeq (Tillich et al., 2017), with tRNAscan-SE v2.0.7 (Chan et al., 2021) to locate tRNAs; we used the available mitogenomes of *Pseudoniphargus stocki* and *Pseudoniphargus carpalis* (Stokkan et al., 2018) as "3rd Party References". A second annotation was conducted using MITOS2 (Bernt et al., 2013), specifying the invertebrate mitochondrial genetic code 5 and the reference dataset RefSeq63 Metazoa.

Considering that no *Niphargus* genome is available in GenBank, we manually checked and refined all annotations, including the possible presence of stop codons of the protein-coding genes and the predicted secondary structure of tRNA genes (using the RNAfold tool available on the ViennaRNA web service; Gruber et al., 2015). To detect initial and terminal codons (stop codon or part of it, such as T– or TA–), sequences were aligned with reference sequences of a very closely related genus (*Pseudoniphargus*: Weber et al., 2021), considering that their sequences can overlap with tRNA sequences Stokkan et al., 2016. The resulting annotated sequences are available in GenBank (accession numbers PV534570, PV534571 and PV534572).

The program mtSVG (available at https://github.com/odethier-ulb/mtSVG) was used to create a visual summary of the final annotated mitogenome of *N. dolenianensis* based on the three mitogenome sequences obtained in the present study. Read coverage depth graphs were created from BAM files using Grace-5.1.17 (Vaught, 1996). Nucleotide diversity analyses of the 13 protein-coding genes and two ribosomal RNA genes were conducted using the packages pegas (Paradis, 2010) and seqinR (Charif and Lobry, 2007) in RStudio 4.4.1.

### Phylogenetic reconstruction

Together with our three Niphargidae mitogenome sequences, we included a selection of representative complete mitogenomes from the amphipod families Pseudoniphargidae (Stokkan et al., 2018), Gammaridae (Cormier et al., 2018; Macher et al., 2017; Mamos et al., 2021), Metacran-gonyctidae (Bauzà-Ribot et al., 2009, 2012), Crangonyctidae (Benito et al., 2021), Talitridae (Kumar Patra et al., 2019), and Hyalellidae, as well as two isopod species (Kilpert et al., 2012; Kilpert and Podsiadlowski, 2006), as an outgroup (all GenBank accession numbers available in Table 2. Families were selected based on their putative affinity with Niphargidae following (Copilaş-Ciocianu et al., 2020). Unfortunately, the mitogenome sequences for amphipods are limited to a few families; the goal of tree building is to allocate the niphargids in an updated, mitogenome-based phylogeny to identify its sister family. The translated protein sequences of all protein-coding genes were concatenated and used to build a maximum-likelihood (ML) phylogenetic tree using the program IQtree2 (Nguyen et al., 2015) with the invertebrate mitochondrial amino acid substitution model (MtInv) (Le et al., 2017), and node support was assessed using 50,000 ultrafast bootstrap replicates (Hoang et al., 2018). The evolutionary model employed (MtInv) is the usual one implemented in IQ-TREE for amino acids and the same used by Macher et al., 2023 for the purpose of comparison. Following Macher et al., 2023, protein-coding genes evolve in a more predictable manner than erratic ribosomal units and can be confidently aligned; therefore, rDNA genes (*rrnL* and *rrnS*) were not included in the analysis. A linear representation of the order of protein-coding genes in each mitogenome was generated using the program mtSVG and these images were added next to the leaves of the ML phylogenetic tree.

**Table 2.**
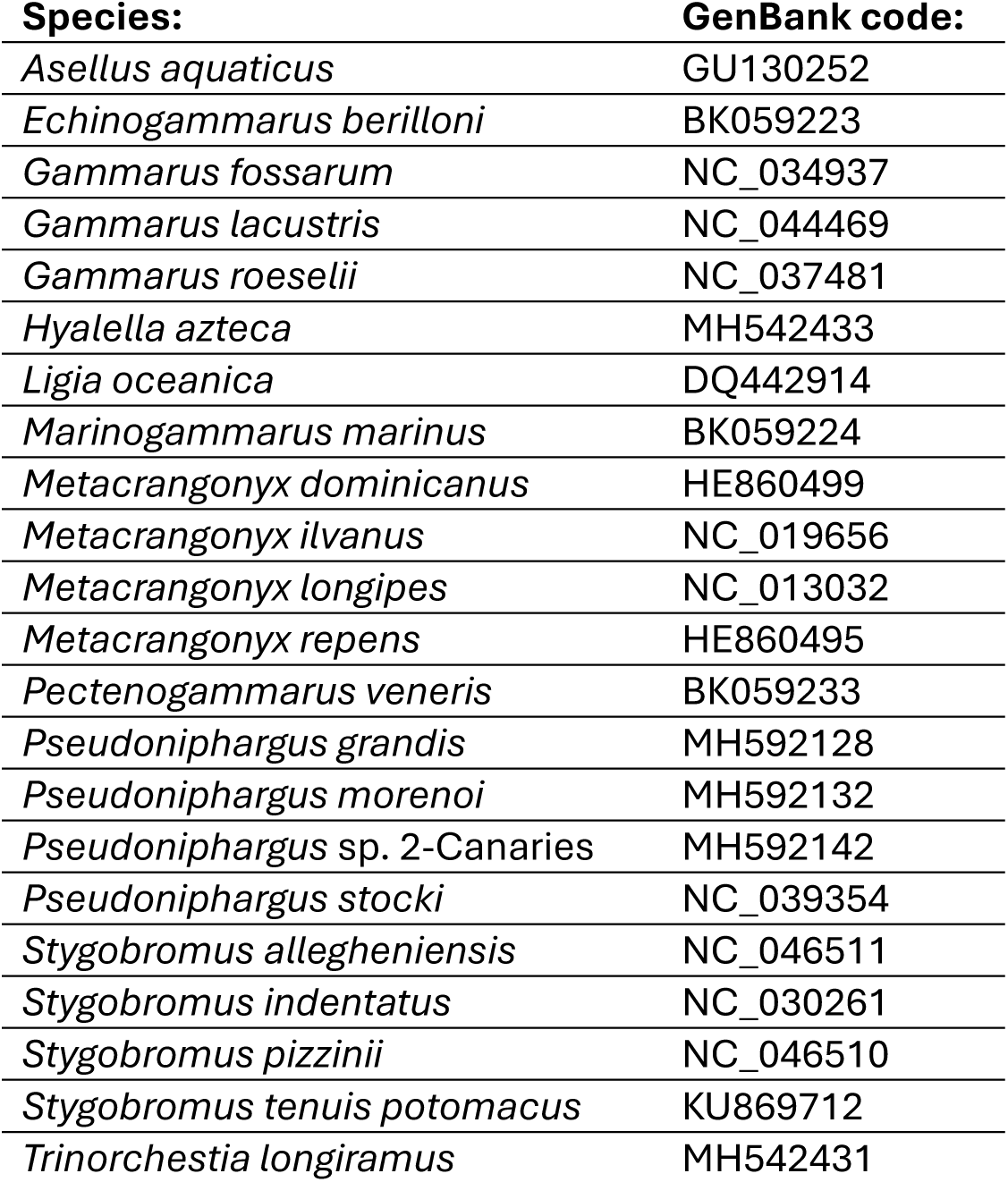
List of all the complete mitochondrial sequences downloaded from GenBank used for mitochondrial genome analysis and integrated with new ones used to build the ML tree.

## Results

The final mitochondrial genome sequences of the three *Niphargus dolenianensis* individuals ranged from 14,964 to 15,097 bp in length (Fig.2). The details of their annotation are reported in Table 3. Their global GC content was 24.6% for FS_24.009, 23.5% for FS_24.010 and 23.0% for FS_24.011.

**Figure 2.**
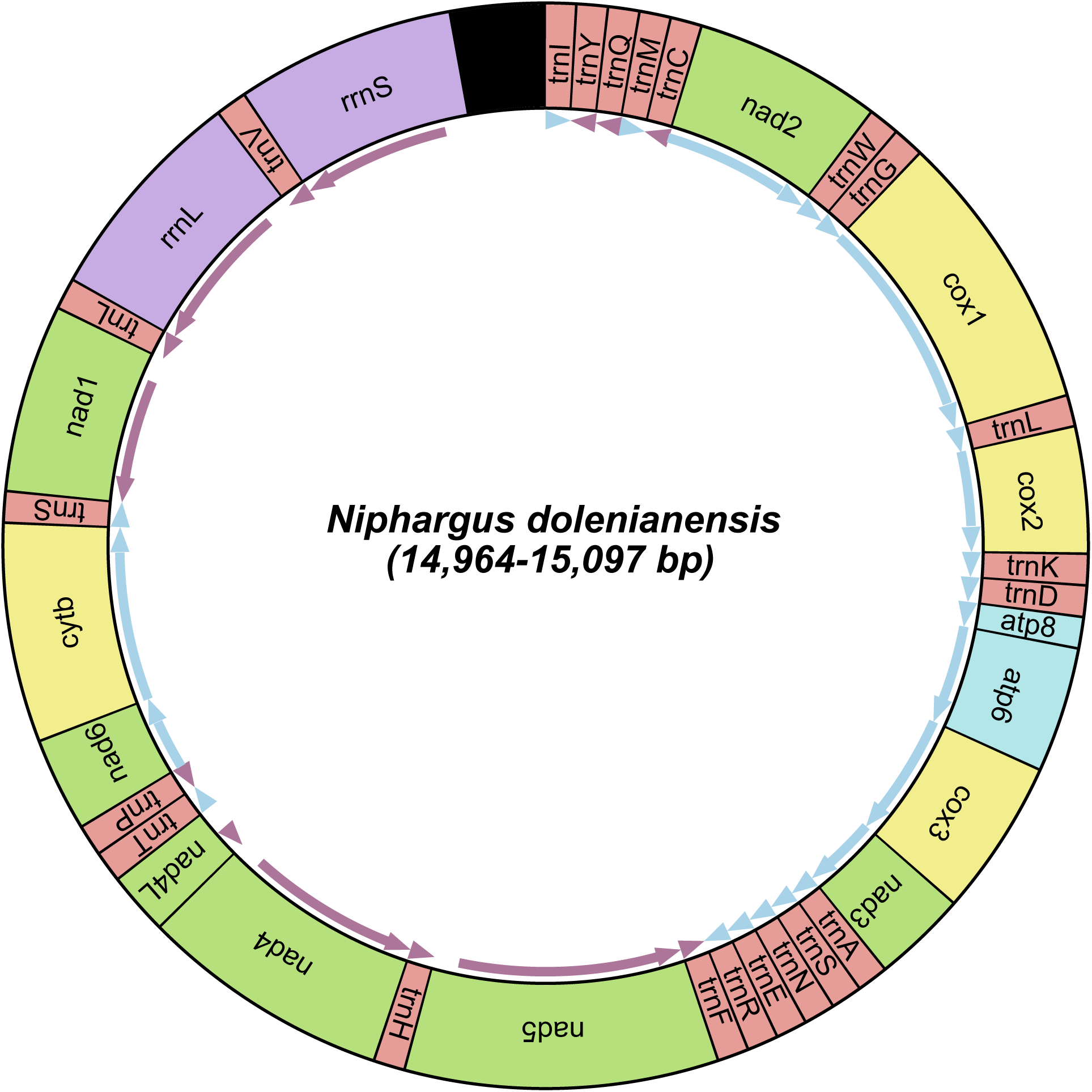
Organization of the mitochondrial genome of *Niphargus dolenianensis* showing the arrangement of the 13 protein-coding genes, the twelve tRNAs, the two rRNA genes and the putative control region (in black).

**Table 3.** Position of all coding and non-coding genes, number of aminoacids in PCGs and their start and stop codons. The underlined numbers represent the position where genes overlap.

| Region | FS_24.009 |  |  |  | FS_24.010 |  |  |  | FS_24.011 |  |  |  |
| --- | --- | --- | --- | --- | --- | --- | --- | --- | --- | --- | --- | --- |
|  | Position | AA | Start | Stop | Position | AA | Start | Stop | Position | AA | Start | Stop |
| *trnI* | 1-61 |  |  |  | 1-61 |  |  |  | 1-60 |  |  |  |
| *trnY* | 64-<u>122</u> |  |  |  | 64-<u>122</u> |  |  |  | 113-<u>171</u> |  |  |  |
| *trnQ* | <u>119</u>-176 |  |  |  | <u>119</u>-176 |  |  |  | <u>168</u>-225 |  |  |  |
| *trnM* | 227-287 |  |  |  | 274-334 |  |  |  | 277-337 |  |  |  |
| *trnC* | 288-346 |  |  |  | 335-393 |  |  |  | 338-396 |  |  |  |
| ***nad2*** | **348-1338** | **330** | **ATT** | **T--** | **395-1385** | **330** | **ATA** | **T--** | **398-1388** | **330** | **ATT** | **T--** |
| *trnW* | 1339-1401 |  |  |  | 1386-1448 |  |  |  | 1389-1451 |  |  |  |
| *trnG* | 1406-1469 |  |  |  | 1452-1515 |  |  |  | 1455-1518 |  |  |  |
| ***cox1*** | **1470-3006** | **512** | **ATA** | **T--** | **1516-3052** | **512** | **ATA** | **T--** | **1519-3055** | **511** | **ATT** | **T--** |
| *trnL2* | 3007-3068 |  |  |  | 3053-3114 |  |  |  | 3056-3117 |  |  |  |
| ***cox2*** | **3069-3741** | **224** | **ATA** | **T--** | **3115-3787** | **224** | **ATA** | **T--** | **3118-3790** | **224** | **ATA** | **T--** |
| *trnK* | 3742-3802 |  |  |  | 3788-3848 |  |  |  | 3791-3851 |  |  |  |
| *trnD* | 3803-3862 |  |  |  | 3849-3908 |  |  |  | 3852-3911 |  |  |  |
| ***atp8*** | **3863-4024** | **53** | **ATT** | **TAA** | **3909-4070** | **53** | **ATT** | **TAA** | **3912-4073** | **53** | **ATC** | **TAA** |
| ***atp6*** | **4018-4686** | **222** | **ATG** | **TAA** | **4064-4732** | **222** | **ATG** | **TAA** | **4067-4735** | **222** | **ATG** | **TAA** |
| ***cox3*** | **4687-5475** | **262** | **ATG** | **TAA** | **4733-5524** | **263** | **ATG** | **TAA** | **4736-5524** | **262** | **ATG** | **TAA** |
| ***nad3*** | **5484-<u>5837</u>** | **117** | **ATG** | **TAG** | **5530-<u>5883</u>** | **117** | **ATG** | **TAG** | **5533-<u>5886</u>** | **117** | **ATG** | **TAG** |
| *trnA* | 5836-5896 |  |  |  | 5882-5942 |  |  |  | 5885-5945 |  |  |  |
| *trnS1* | <u>5896</u>-<u>5947</u> |  |  |  | <u>5942</u>-<u>5993</u> |  |  |  | <u>5945</u>-<u>5996</u> |  |  |  |
| *trnN* | <u>5947</u>-6008 |  |  |  | <u>5993</u>-6054 |  |  |  | <u>5996</u>-6057 |  |  |  |
| *trnE* | <u>6006</u>-<u>6069</u> |  |  |  | <u>6052</u>-<u>6115</u> |  |  |  | <u>6055</u>-<u>6118</u> |  |  |  |
| *trnR* | 6064-6122 |  |  |  | 6110-6168 |  |  |  | 6113-6171 |  |  |  |
| *trnF* | <u>6121</u>-6180 |  |  |  | <u>6167</u>-6225 |  |  |  | <u>6170</u>-6229 |  |  |  |
| ***nad5*** | **6181-7888** | **569** | **ATA** | **T--** | **6226-7933** | **569** | **ATA** | **T--** | **6230-7937** | **569** | **ATA** | **T--** |
| *trnH* | 7889-7948 |  |  |  | 7934-7993 |  |  |  | 7938-7997 |  |  |  |
| ***nad4*** | **7949-9263** | **438** | **ATG** | **T--** | **7994-9308** | **439** | **ATG** | **T--** | **7998-9312** | **438** | **ATG** | **T--** |
| ***nad4L*** | **9257-9547** | **96** | **ATG** | **TAA** | **9302-9592** | **96** | **ATG** | **TAA** | **9306-9596** | **96** | **ATG** | **TAA** |
| *trnT* | 9551-<u>9610</u> |  |  |  | 9595-<u>9654</u> |  |  |  | 9599-<u>9658</u> |  |  |  |
| *trnP* | <u>9609</u>-9670 |  |  |  | <u>9653</u>-9714 |  |  |  | <u>9657</u>-9718 |  |  |  |
| ***nad6*** | **9673-<u>10170</u>** | **165** | **ATG** | **TAA** | **9729-<u>10214</u>** | **161** | **ATT** | **TAA** | **9733-<u>10218</u>** | **161** | **ATT** | **TAA** |
| ***cob*** | **<u>10170</u>-<u>11309</u>** | **379** | **ATG** | **TAA** | **<u>10214</u>-<u>11353</u>** | **379** | **ATG** | **TAA** | **<u>10218</u>-<u>11357</u>** | **379** | **ATG** | **TAA** |
| *trnS2* | <u>11308</u>-<u>11367</u> |  |  |  | <u>11352</u>-<u>11411</u> |  |  |  | <u>11356</u>-<u>11415</u> |  |  |  |
| ***nad1*** | **<u>11365</u>-<u>12276</u>** | **303** | **ATT** | **TAA** | **<u>11409</u>-<u>12320</u>** | **303** | **ATT** | **TAA** | **<u>11413</u>-<u>12324</u>** | **303** | **ATT** | **TAA** |
| *trnL1* | 12286-<u>12347</u> |  |  |  | 12330-<u>12391</u> |  |  |  | 12334-<u>12395</u> |  |  |  |
| *rrnL* | <u>12325</u>-13359 |  |  |  | <u>12369</u>-13404 |  |  |  | <u>12373</u>-13408 |  |  |  |
| *trnV* | 13384-13429 |  |  |  | 13428-13473 |  |  |  | 13433-13478 |  |  |  |
| *rrnS* | 13430-14487 |  |  |  | 13474-14528 |  |  |  | 13479-14531 |  |  |  |
| CR | 14488-15097 |  |  |  | 14529-14964 |  |  |  | 14532-15069 |  |  |  |

For the sample FS_24.009 obtained from Illumina and nanopore, respectively, 2.82 and 1.71 Gbp, for FS_24.010, 2.66 and 0.69 Gbp, and for FS_24.011, 0.95 and 1.70 Gbp. The coverage depth obtained for the assembled mitogenomes generated in NovaSeq 6000 Illumina short reads ranged from approximately 40 to 250X; similarly, the coverage depth for nanopore long reads ranged from 40 to 300X (Fig.3). nanopore coverage depth profiles were less even than those for Illumina, possibly because of biases caused by the PCR amplification step in the Rapid PCR Barcoding Kit protocol.

**Figure 3.**
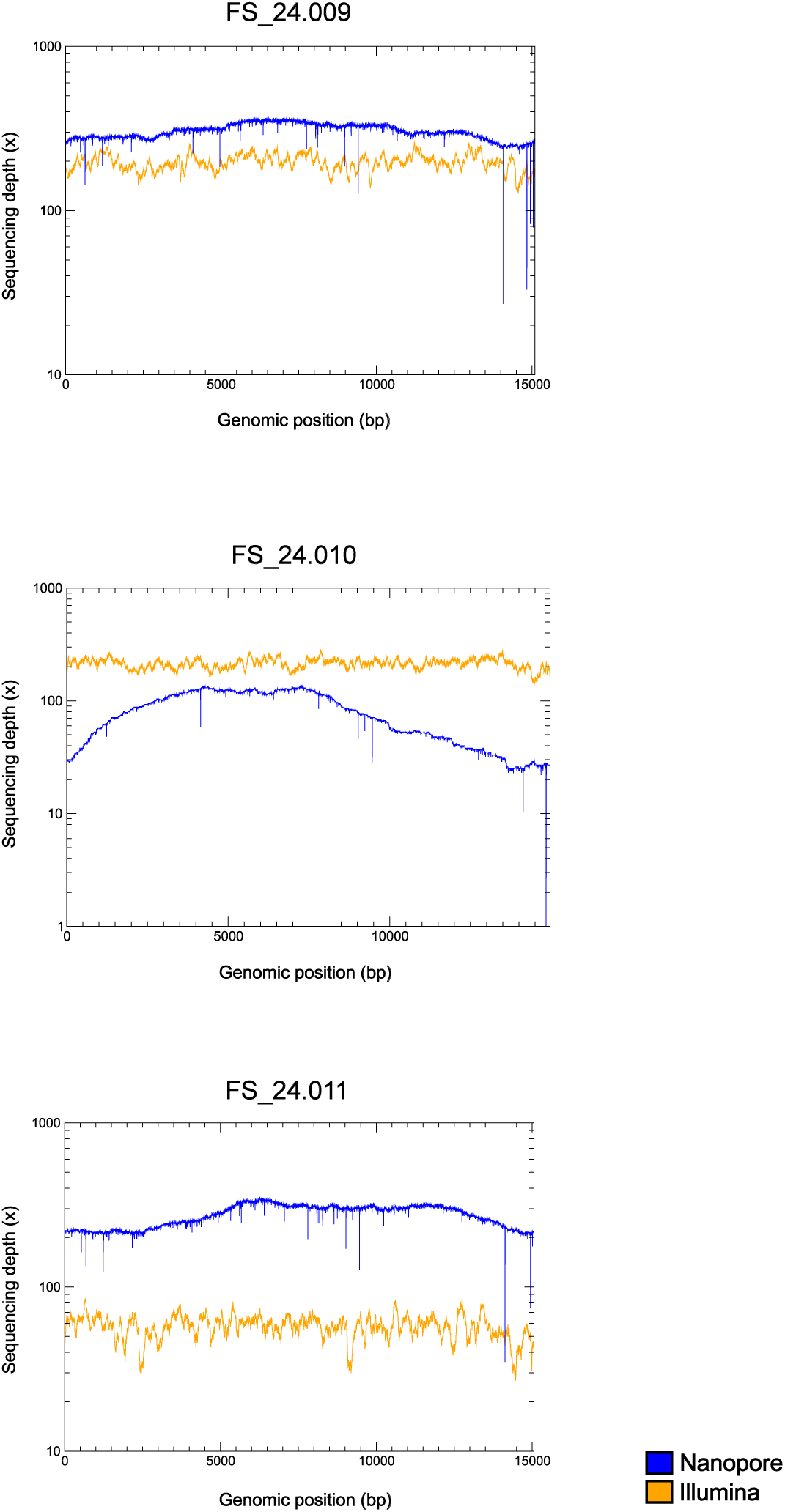
Read coverage graph comparing Illumina and nanopore data for the three *Niphargus* individual. Graphs with logarithmic y-axis.

The predicted secondary structures of the tRNA genes are shown in Fig.4. Only tRNA-Gly and tRNA-Leu2 were identical across all three individuals, whereas all other tRNA sequences had a few base differences (highlighted in pink in Fig.4). These differences were mostly substitutions (most often located outside of stem structures, except for a few mutations in stems that either did not disturb base pairing or, in the case of tRNA-Met, that was compensated by a mutation in the facing position). Of the 22 tRNAs annotated in the mitochondrial genome of *Niphargus dolenianensis*, 19 had the typical 3-arm structure, whereas tRNA-Ser1 and tRNA-Val lacked the dihydrouridine (DHU) arm and tRNA-Phe lacked the T-arm. The structure of tRNA-Val of FS_24.011 could not be predicted *de novo* from its sequence using the Vienna server because of the extra G-A pairing of this sequence compared with the other two. Therefore, we constrained its folding to the structure shown in Fig.4 using the results obtained from the two other individuals.

**Figure 4.**
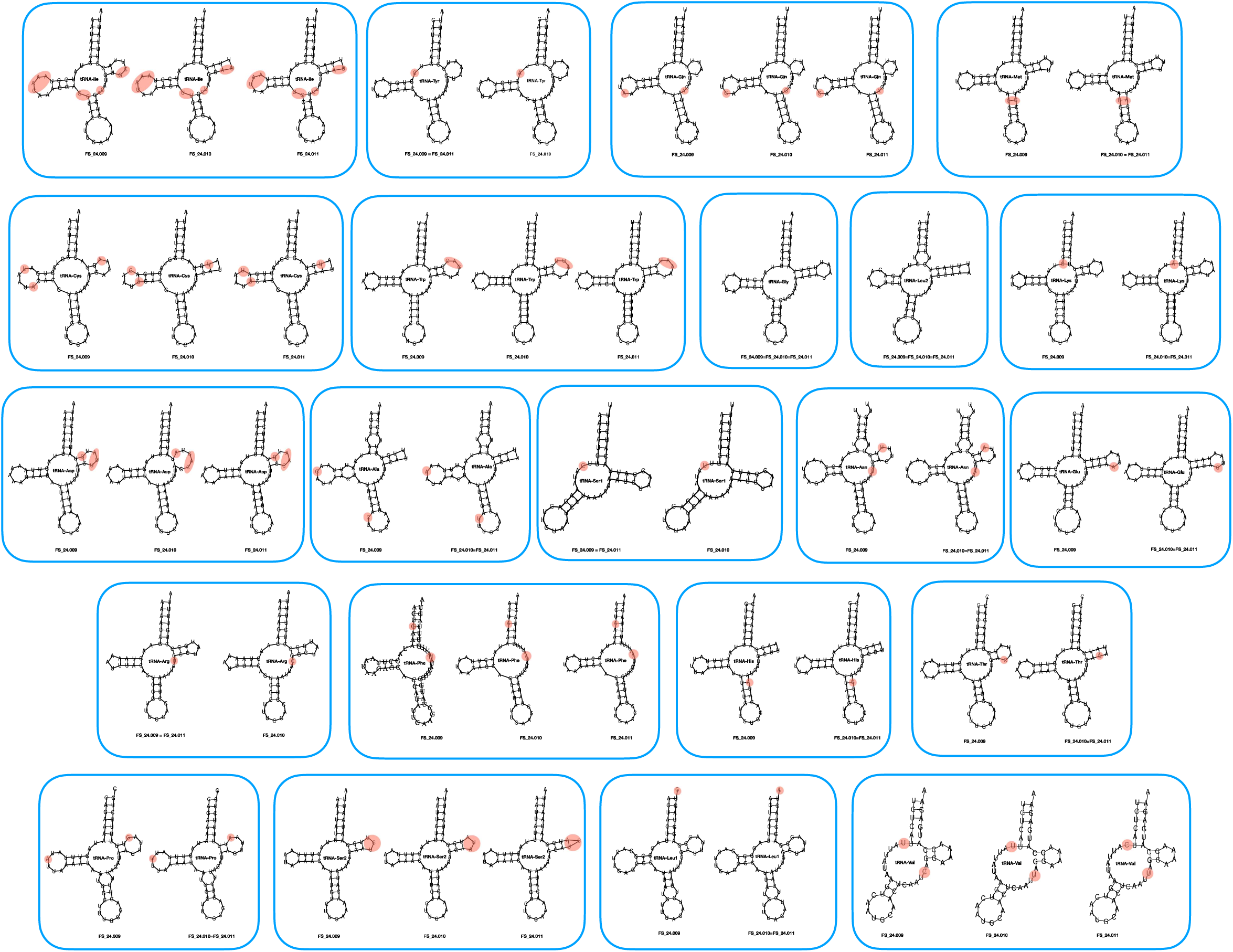
Predicted secondary structure of the 22 tRNA genes.

The nucleotide diversity (*π*) values for each protein-coding and ribosomal RNA gene are shown in Fig.5. The values range from 0.06 to 0.09 for the protein-coding genes, whereas it is only 0.03-0.04 for the two ribosomal RNA genes.

**Figure 5.**
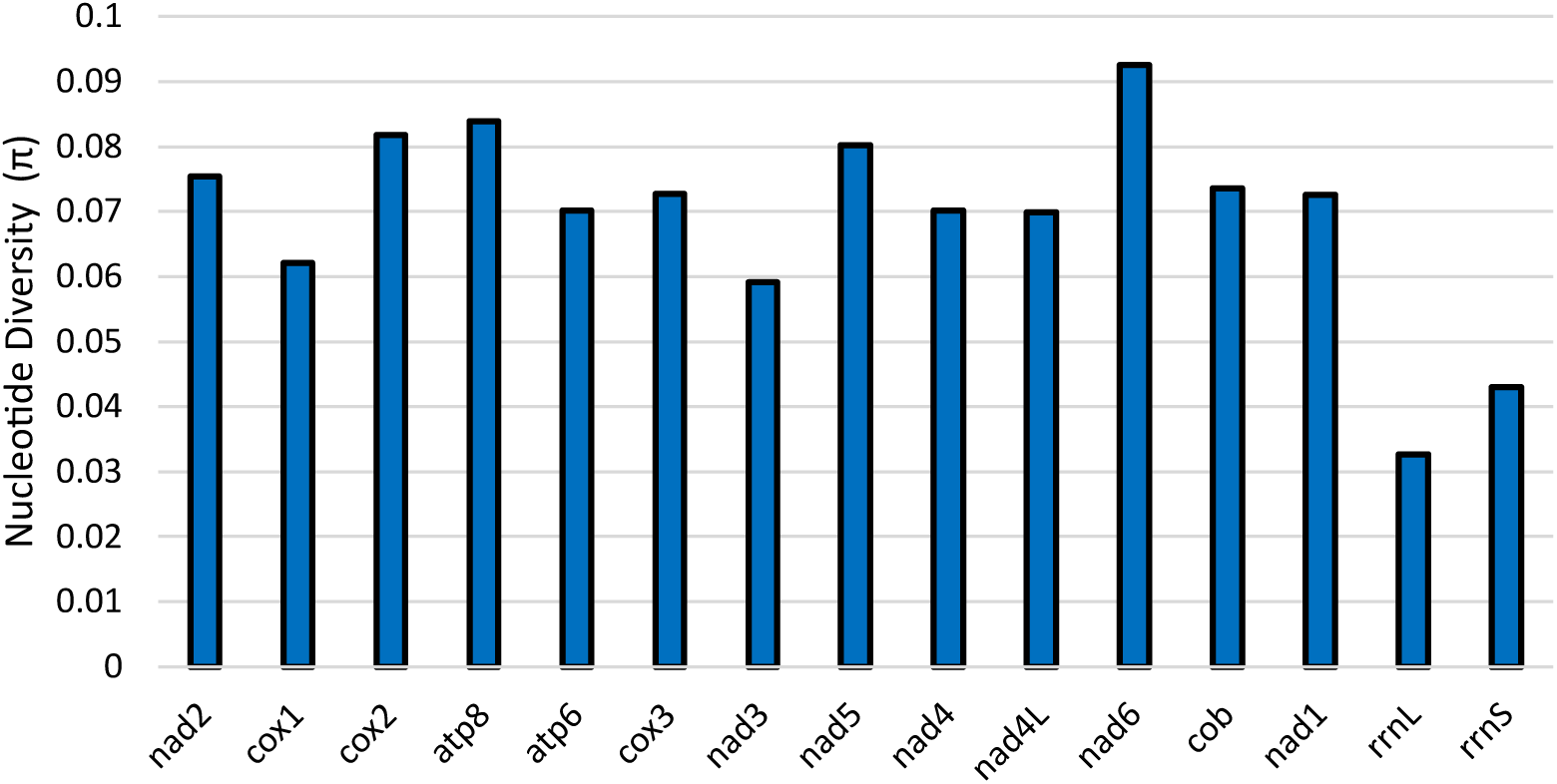
Nucleotide diversity of the three *Niphargus* mitogenomes for protein-coding genes and rRNA genes.

There were no significant differences between the Flye and Hifiasm mitogenome assemblies obtained from the nanopore long reads: for FS_24.009, the two were perfectly identical, whereas in the case of FS_24.010 and FS_24.011, there were only two differences between the Flye and Hifiasm mitogenome assemblies, in both short indels at the level of long homopolymers. These indels and several others (always located in long homopolymers) in the nanopore-assembled sequences were corrected through polishing using Illumina reads. In the mitogenome of FS_24.009, the differences between the Flye nanopore assembly before and after Illumina polishing were: a one-base indel in *nad4L* and a three-base indel in *rrnS*; in the mitogenome of FS_24.010, a one-base indel in *atp6*, another one in *nad4L*, and a two-base indel in *rrnS*; in the mitogenome of FS_24.011, a one-base indel in *nad4L*, and a two-base indel in *rrnS*.

By contrast, the Illumina-only assemblies obtained using NOVOplasty had many discrepancies with the nanopore-assembled, Illumina-polished accurate sequences: the majority of the differences were detected in the control regions with various insertions, and few indels and base substitutions were also found in tRNAs as well as in some protein-coding genes; for the individual FS_24.009, we detected four substitutions in trnY and one substitution in *nad3* and *nad4L*; for the individual FS_24.010, we detected four substitutions in trnY, a 43 bases deletion between trnQ and trnM, and one base substitution in *nad3*; for the individual FS_24.011, we detected four substitutions in trnY, a 55 bases deletion between trnI and trnY, a 5 bases deletion between trnG, and one base substitution in *nad3*.

The ML tree obtained from this analysis is shown in Fig.6. The newly annotated mitogenome of *Niphargus* formed a clade with *Pseudoniphargus*, supported by a 99% ultrafast bootstrap value. No difference was found in the protein-coding gene order between Niphargidae and Pseudoniphargidae, but some differences were detected between more distant amphipod families. Most Gammaridae and all Crangonyctidae had the same gene order as Niphargidae and Pseudoniphargidae, with the exception of *Marinogammarus marinus* for which *nad5*, *rrnS* and *rrnL* were translocated. In the case of Metacrangonyctidae, *cytb* changed position and orientation; whereas in Hyalellidae, *nad1* changed orientation but remained in the same location. The talitrid *Trinorchestia longiramus* had *nad3* located before *cox1* and its *cytb* and *nad6* were swapped.

**Figure 6.**
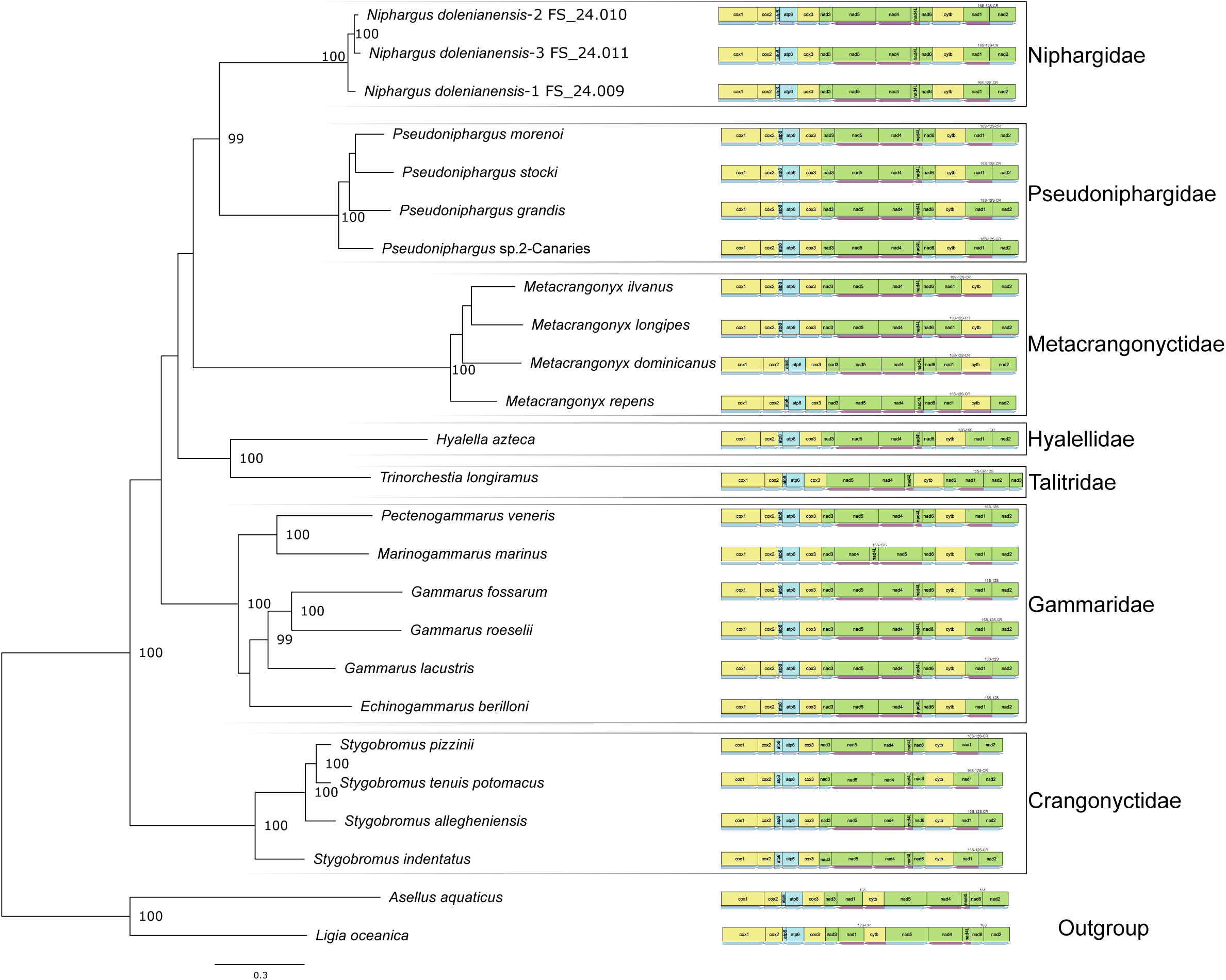
Maximum likelihood phylogenetic tree obtained from translated protein sequences of all mitochondrial protein-coding genes. Different families are indicated by black lines. Labels of nodes with a probability below 95 % were removed. 16S, 12S and CR represent the position of the *rrnL* and *rrnS* genes as well as the control region, when available from the annotation on Genbank.

## Discussion

Like other metazoans, the mitochondrial genome of crustaceans typically consists of a circular double-stranded DNA molecule ranging from 12 to 20 kb, with highly conserved gene content. It contains 13 protein-coding genes (PCGs), two ribosomal RNA genes (rRNAs), 22 transfer RNA genes (tRNAs), and a large non-coding region where replication of the mitochondrial genome is initiated, called the D-loop or control region (CR) (Boore, 1999). In this study, we present the results obtained from the first three complete mitogenomes of the genus *Niphargus* belonging to the species *N. dolenianensis*.

Nanopore assembly followed by careful Illumina polishing (first automatically, then manually) yielded genome annotations that were highly consistent with published amphipod mitogenomes, particularly with those of Pseudoniphargidae, which, based on our phylogenetic reconstruction using protein-coding genes, is the sister family to Niphargidae, as already hypothesized by (Weber et al., 2021). Most mt-tRNAs fold into the same cloverleaf secondary structure as nuclear-encoded tRNA sequences, comprising four stems and three loops (Jühling et al., 2012). However, as reported in the genus *Pseudoniphargus* Stokkan et al., 2016, the tRNA-Ser1 and tRNA-Val of *Niphargus dolenianensis* lacked the dihydrouridine (DHU) arm, also common in nearly all metazoans (Ki et al., 2010), while its tRNA-Phe lacked the T-arm.

The tRNA secondary structures generally contained wobble base pairs (G–U, I–U, I–A, and I–C); however, we also noticed some non-canonical paring (e.g. U–U, G–G, U–C, A–A), as frequently observed in other species (Ki et al., 2010).

Nucleotide diversity values were higher for protein-coding genes than for ribosomal RNA genes. Among the protein-coding genes, a higher diversity value was observed for *nad6*, whereas *cox1* had a comparatively lower value, even though the latter is commonly used to identify arthropod species.

Using nanopore for genome skimming allowed us to exclude nuclear mitochondrial DNA segments (numts) because the amplification step during library preparation did not involve any gene-specific primers. The absence of targeted amplification is a key advantage of this technique. Illumina-only assemblies were imperfect, with many structural errors (mainly in the control region and in ribosomal RNA genes, although adjacent tRNA regions were also affected), probably due to short-read assembly, whereas nanopore-only assemblies were plagued by one-based artifactual indels in coding regions (caused by homopolymers longer than 10-12 bases), resulting in reading-frame shifts. However, despite Illumina sequencing providing a higher amount of data (Gbp) per sample (except for FS_24.011, for which the nanopore-only assembly generated more data), this increase did not translate into improvements in either the final assembly quality or coverage depth compared to the nanopore-only assembly.

The fact that the nanopore-only assembly contained some artifactual indels was unexpected, given that nanopore R10.4.1 data have been reported to yield error-free bacterial genome sequences that do not require Illumina polishing (Sereika et al., 2022). As mitochondria are alphaproteobacteria (Fan et al., 2020), the same approach would have been expected to work on mitogenomes as well. However, the mitogenome of *Niphargus dolenianensis* contains many long stretches of polyA and polyT (up to a length of 25 identical nucleotides in a row) that are not typically observed in bacterial genomes and are responsible for the artifactual indels observed. These stretches were mainly observed in the two rDNA genes and in the control region, although in some cases, this kind of error may disrupt protein-coding genes. These errors were manually corrected after checking amino acid translation and comparing Illumina outputs, as well as nanopore-based sequences of the three different, conspecific individuals used in this study. This is most probably caused by the extremely low GC content of the mitochondrial genome of *Niphargus dolenianensis*, which, being lower than 25% GC, falls outside the range of GC content of bacteria and archaea, except for a very few cases such as *Carsonella* (Mann and Chen, 2010). In such cases, both nanopore and Illumina data are required to obtain a highly accurate, error-free mitogenome sequence. This is particularly important when producing the first reference mitogenome sequence for a previously unexplored genus or family, as was the case here. Subsequently, we can conclude that resequencing of other species closely related to the one for which a reference sequence was generated could rely on nanopore only, since all the artifactual errors in the resulting genome assembly will be located at the level of long homopolymers and will be easily detected and corrected by hand. In terms of both cost and turnaround time, nanopore sequencing is markedly faster and more economical, with the potential to generate final raw data in less than two days from the initial processing of animal tissue. Sequencing the mitogenomes of three closely related specimens, as we did, allowed us to cross-compare and refine our gene annotations and was also instrumental in obtaining properly folded sequences for all tRNAs, as the tRNA-Val of FS_24.011 could not be folded properly without relying on information from the other two. Therefore, if we had only sequenced FS_24.011, we could have incorrectly concluded that tRNA-Val was absent in *Niphargus dolenianensis*.

The phylogenetic tree of amphipods, using the complete set of protein-coding genes in the mitochondrial genome, yielded a well-resolved phylogeny and confirmed the close evolutionary relationship between Niphargidae and Pseudoniphargidae. Their clade had very high bootstrap support in its ancestral node. This highlights the usefulness of complete mitochondrial genome sequences for conducting studies at a deeper phylogenetic level (Bauzà-Ribot et al., 2009; Sun et al., 2018), with the known limitations of mitochondrial genomes, mainly due to saturation problems, in resolving some ancient splits that are millions of years old (Phillips et al., 2013). It is well known that the saturation problem of the *cox1* gene in amphipods (Stoch et al., 2024a, 2022) leads to poor resolution of ancient splits. The phylogenetic tree generated using complete mitogenome sequences of amphipods made it possible to resolve some of these basal nodes (Bauzà-Ribot et al., 2013; Höpel et al., 2022; Stokkan et al., 2018). Our research offers valuable insights into nanopore sequencing that can be employed to obtain more precise mtDNA genome sequences, which is promising for gaining a clearer understanding of the evolution and diversification of amphipods.

## Acknowledgements

The authors wish to thank Laurent Grumiau and Florence Rodriguez Gaudray for their assistance with the laboratory work.

AS’s Ph.D. was supported by a Université libre de Bruxelles (ULB) seed grant and by DarCo (The vertical dimension of conservation: a cost-effective plan to incorporate subterranean ecosystems in post-2020 biodiversity and climate change agendas, BIODIV21_0006).

NL’s Ph.D. was supported by a seed grant for the ULB-VUB collaborative research.

FS and JFF were supported by Projet de Recherches’ grant no. T.0078.23 to JFF.

This study is part of the DarCo project funded by Biodiversa+, the European Biodiversity Partnership under the 2021–2022 BiodivProtect joint call for research proposals, co-funded by the European Commission (GA no. 101052342) and the Ministry of Universities and Research (Italy), Agencia Estatal de Investigación—Fundación Biodiversidad (Spain), Fundo Regional para a Ciência e Tecnologia (Portugal), Suomen Akatemia—Ministry of the Environment (Finland), Belgian Science Policy Office (Belgium), Agence Nationale de la Recherche (France), Deutsche Forschungsgemeinschaft e.V. (Germany), Schweizerischer Nationalfonds (grant no. 31BD30_209583, Switzerland), Fonds zur Förderung der Wissenschaftlichen Forschung (Austria), Ministry of Higher Education, Science and Innovation (Slovenia), and the Executive Agency for Higher Education, Research, Development and Innovation Funding (Romania).

We sincerely thank the anonymous referees and two anonymous reviewers for their constructive comments and valuable suggestions, which greatly improved the quality of this manuscript.

## Author contributions

AS conducted most of the laboratory work, contributed to data analysis and interpretation, and drafted the manuscript with input from all authors. NL contributed to the laboratory work. FS collected all samples and contributed to data analysis, mitogenome annotation, and interpretation. JFF conducted most of the data analyses and contributed to their interpretation and validation. All authors reviewed and approved the final manuscript.

## Command lines

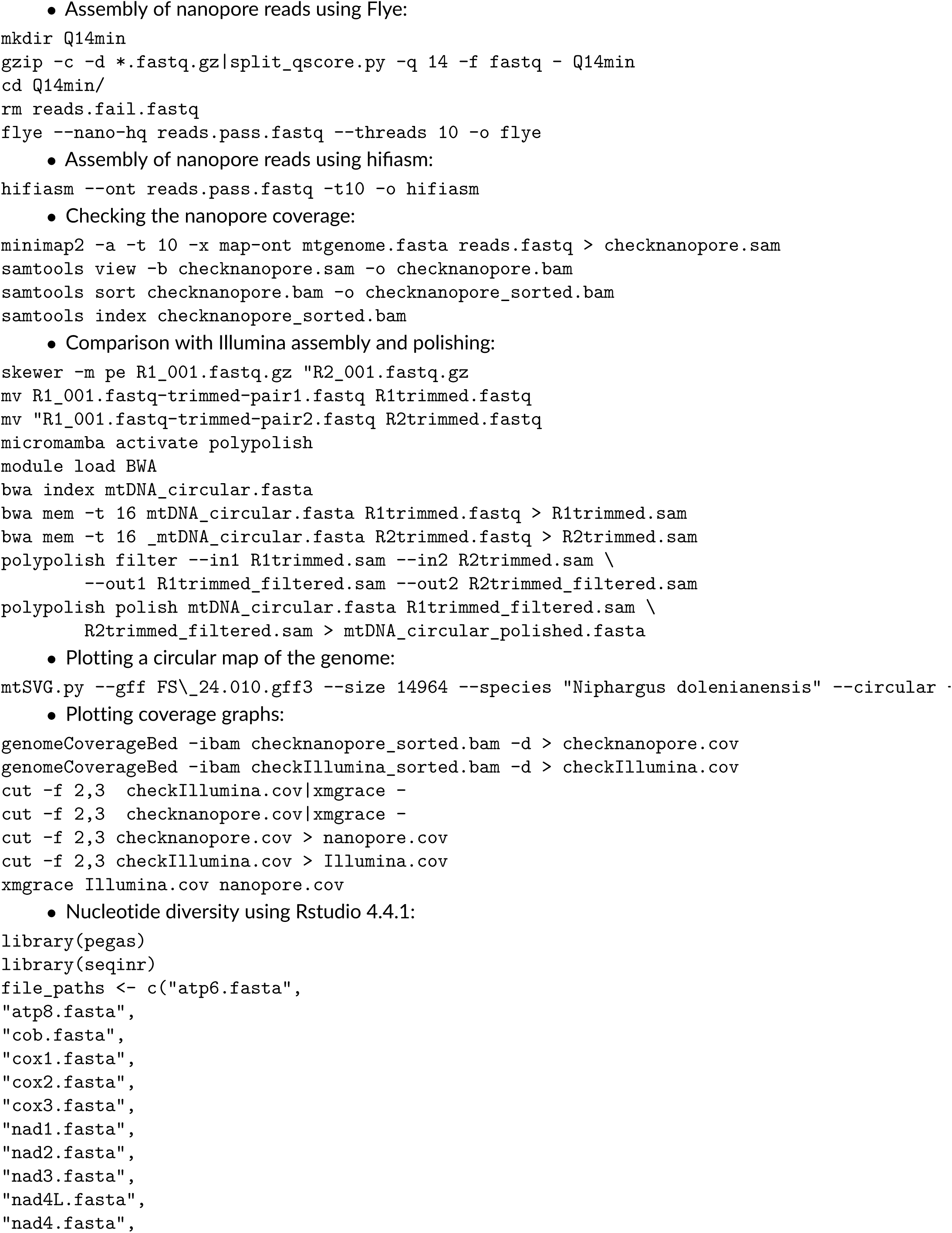
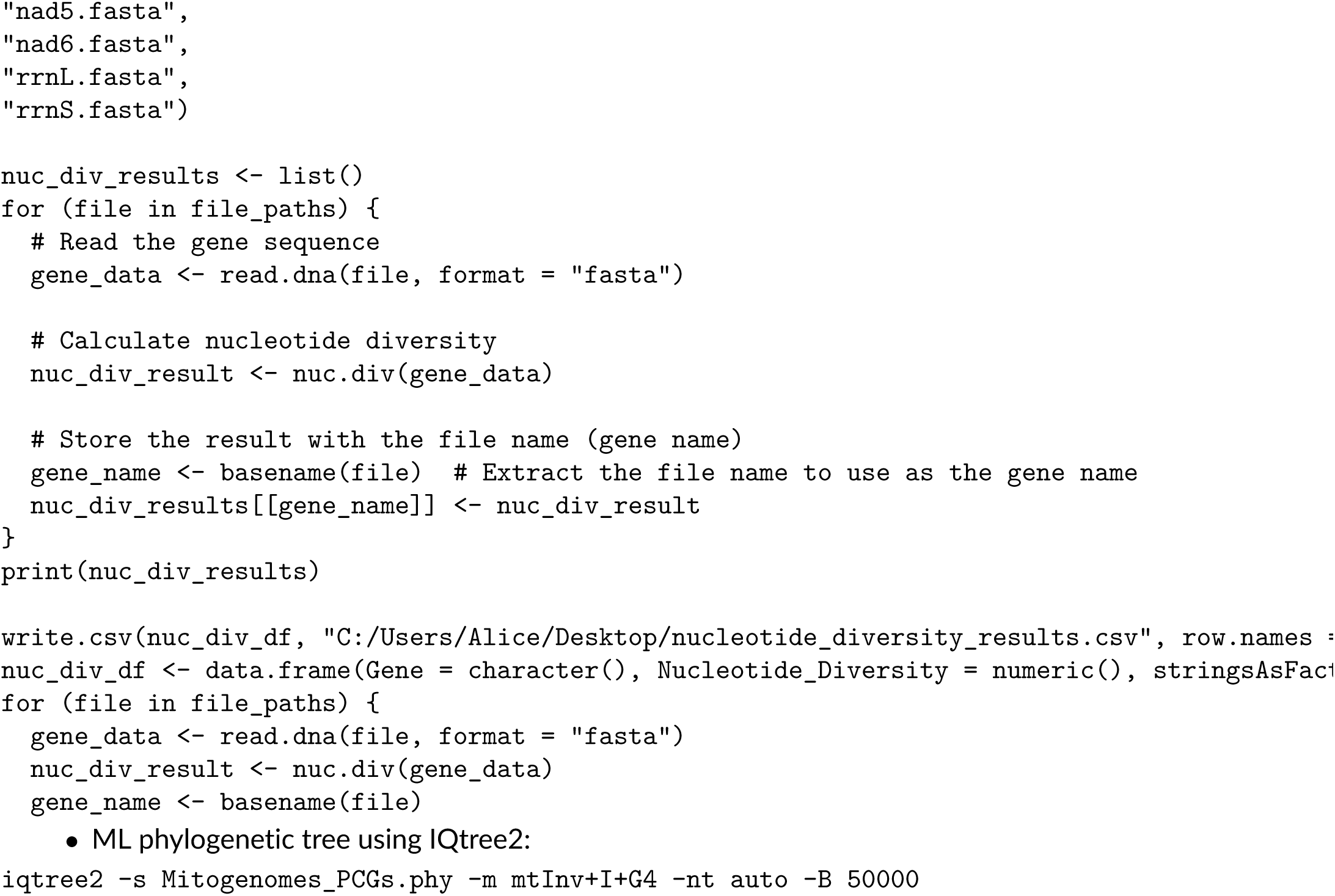

